# Enhancing proteoform sequence coverage using top-down mass spectrometry with in-source fragmentation and middle-down mass spectrometry

**DOI:** 10.1101/2025.09.26.678817

**Authors:** Xingzhao Xiong, Letu Qingge, Binhai Zhu, Xiaowen Liu

**Affiliations:** Deming Department of Medicine, School of Medicine, Tulane University, New Orleans, LA, USA; Department of Computer Science, North Carolina A&T State University, Greensboro, North Carolina, USA; Gianforte School of Computing, Montana State University, Bozeman, Montana, USA

## Abstract

The study of complex proteoforms with mutations and post-translational modifications has gained increasing attention with the advancement of mass spectrometry (MS)-based techniques. Achieving high proteoform sequence coverage by MS is essential for accurately characterizing these complex proteoforms. Extensive efforts have been made to increase proteoform sequence coverage using deep bottom-up and top-down MS strategies. In this study, we evaluated top-down and middle-down MS approaches for enhancing proteoform sequence coverage using three proteins: ubiquitin, myoglobin, and carbonic anhydrase II. In the top-down MS approach, we applied in-source fragmentation (ISF) to generate pseudo-MS^3^ spectra, thereby improving sequence coverage. For the middle-down MS strategy, we performed short-duration enzymatic digestions to produce longer peptides that preserve more proteoform sequence information. Our experimental results demonstrated that ISF and partial digestion significantly increased the sequence coverage of the proteins, achieving coverage greater than 90%.

## 1 Introduction

In mass spectrometry (MS)-based proteomics, high proteoform sequence coverage is essential to proteoform characterization, in which all post-translational modifications (PTMs) on proteoforms need to be identified and characterized [1]. Achieving almost 100% sequence coverage is especially important to study complex proteoforms with multiple PTM sites, such as histone proteoforms and phosphorylated ones [2-4]. *De novo* sequencing of proteoforms [5], such as antibodies, also demands almost 100% sequence coverage. Because of this, many efforts have been made to increase proteoform coverage using MS.

Bottom-up, middle-down, and top-down MS approaches have complementary strengths in increasing proteoform coverage [6-8]. In bottom-up proteomics (BUP), proteins are digested using trypsin or other proteases to produce short peptides [9-11], which are subsequently identified using tandem mass spectrometry (MS/MS) coupled with liquid chromatography (LC) [12, 13]. Bottom-up MS offers high fragment ion coverage for peptides generated by enzyme digestion, facilitating the identification and localization of PTMs on the peptides [1, 9]. In addition, combining bottom-up MS data generated by multiple enzymes separately can further increase proteoform sequence coverage for proteoform characterization [14]. However, it is still challenging to achieve high sequence coverage in proteome level studies, in which complex samples with thousands or tens of thousands of proteins are analyzed [1, 15]. In addition, because the combinatorial patterns of PTMs are lost during digestion, it is challenging to characterize whole proteoforms with multiple PTMs when several similar proteoforms co-exist in the sample.

Top-down proteomics (TDP) offers unique advantages for studying complex proteoforms with multiple PTM sites, as it directly analyzes intact proteoforms without prior digestion and preserves the combinatorial information of PTM sites on proteoforms [16-18]. Recent advancements in TDP have made it the method of choice for investigating large proteoforms [19-24]. In TDP, commonly used fragmentation methods, such as higher-energy collisional dissociation (HCD), collision-induced dissociation (CID), and electron-transfer association (ETD), typically target the most labile bonds, leading to limited sequence coverage [16] and making it challenging to confidently localize PTMs [25, 26]. Although numerous efforts have been made to increase proteoform sequence coverage by combining various fragmentation methods and optimizing MS parameter settings [4, 19, 27, 28], top-down MS still often fails to obtain complete sequence coverage of proteoforms.

Middle-down proteomics (MDP) is an alternative approach to BUP and TDP, which analyzes peptides/proteoforms that are longer than those in BUP and shorter than those in TDP [29-32]. The peptides/proteoforms analyzed in MDP are often generated from digestion using enzymes with fewer digestion sites, such as LysC [33]. MDP provides distinct advantages in proteoform characterization. Compared with BUP, MDP retains more combinatorial PTM information within relatively long peptides or proteoforms. In contrast to TDP, MDP achieves higher sequence coverage by analyzing these longer peptides/proteoforms [34].

Here we evaluated two methods for increasing proteoform sequence coverage using three proteins with varying molecular weights: ubiquitin (8.6 kDa), myoglobin (17 kDa), and carbonic anhydrase II (CA2; 29 kDa). The first method was top-down MS with in-source fragmentation (ISF), which fragments intact proteoforms at the ion source and then generates MS/MS spectra of the fragmented proteoforms, resulting in pseudo MS^3^ spectra [35]. The second was middle-down MS with short digestion time, which can generate long peptides/proteoforms [29] with combinatorial PTM information. Our experimental results demonstrated that both methods substantially increased proteoform sequence coverage of the proteins compared with standard top-down MS.

## 2 Methods

### 2.1 Chemicals and materials

Ammonium bicarbonate (ABC), dithiothreitol (DTT), iodoacetamide (IAA), and urea were purchased from Sigma (St. Louis, MO). Acetonitrile (ACN, HPLC grade), formic acid (FA), isopropanol (IPA, HPLC grade), methanol (MeOH, HPLC grade) and water (HPLC grade) were obtained from Thermo Scientific, (Waltham, MA, USA). Standard proteins were purchased from Sigma-Aldrich: ubiquitin (U6253), myoglobin (M0630), and CA2 (C2624). Enzymes were purchased from Sigma-Aldrich: trypsin (T1426), Roche: chymotrypsin (11418467001) and GluC (10791156001), and New England Biolabs: AspN (P8104S) and LysC (P8109S).

### 2.2 Top-down MS

Top-down MS experiments were carried out using an Ultimate 3000 LC system coupled with an Orbitrap Fusion Lumos mass spectrometer (Thermo Scientific, Waltham, MA, USA). Ubiquitin, myoglobin, and CA2 were analyzed separately. A total of 100 μg protein was dissolved in 100 μL of water. Myoglobin and CA2 were reduced with 1 μL of 1 M DTT at 55 °C for 45 minutes, while the reduction step was omitted for ubiquitin, as it does not contain disulfide bonds. The resulting protein was then diluted in a solution containing 50% water, 50% MeOH, and 0.1% FA to the concentration of 20 ng/μL. A total of 10 ng protein was loaded and separated using reverse phase liquid chromatography (RPLC) with a C2 column (300 Å, 3 µm, 100 μm i.d., 60 cm length, CoAnn, Richland, WA). A 30 min gradient was used for protein separation, and the flow rate was 300 μL /min. Mobile phase A was water with 0.1% FA; mobile phase B consisted of 60% ACN, 15% IPA, and 25% water with 0.1% FA. The gradient for phase B was set as follows: 0–2 min, 5% to 35%; 2–5 min, 35% to 50%; and 5–30 min, 50% to 80%.

Full MS scans were collected with a resolution of 120,000 for ubiquitin and 240,000 for myoglobin and CA2 with 3 microscans. The scan range was 720-1200 m/z, and the AGC target was 1×10^6^ with a maximum injection time of 500 ms. A total of 3 CID MS/MS scans were collected for each MS scan using the data dependent acquisition (DDA) mode. MS/MS data were acquired using a resolution of 60,000 with 1 microscan, an isolation window of 1.2 m/z, and a scan range of 400-2000 m/z. The normalized collision energy (NCE) was set to 35%, the AGC target was set to 1× 10^6^, and the maximum injection time was set to 500 ms for ubiquitin and 1000 ms for myoglobin and CA2. We conducted MS runs for each of the three proteins using the MS settings described above, varying the ISF energy from 0 V to 100 V in a 10 V increment (11 settings in total). For each ISF energy setting, experiments were conducted in technical triplicate.

### 2.3 Top-down MS data analysis

Top-down MS data were converted to mzML files using the tool msconvert in ProteoWizard [36]. Then TopFD [37] (version 1.7.9; parameter settings in Supplemental **Table S1**) was used to identify proteoform features and deconvolute spectra in the mzML files. The resulting deconvoluted spectra were searched against the protein database containing only the target protein sequence using TopPIC [38] (version 1.7.9; parameter settings in Supplemental **Table S2**), in which unknown mass shifts were not allowed and water loss on all 20 amino acids was chosen as the variable PTM. For each MS data file, TopFD reported a feature intensity for each proteoform, which is the sum of all peak intensities corresponding to various isotopic compositions, charge states, and retention times observed in MS1 spectra. The proteoform with the highest feature intensity in the MS data with the 0V ISF energy was selected as *the reference proteoform*.

The reference proteoform of ubiquitin was observed in the MS data of all the 11 ISF conditions. But in some MS data files of myoglobin and CA2 with a high ISF energy, the reference proteoform was not identified. For these files, we selected an ISF proteoform as an alternative reference proteoform with a high proteoform abundance (see Results). In an MS data file, the *reference elution time* is defined as the apex retention time of the reference proteoform if the reference proteoform is observed and as the apex retention time of the alternative reference proteoform otherwise.

For each of the three proteins, we searched for fragment proteoforms resulting from ISF of the reference proteoform, referred to as *ISF proteoforms*. To find highly confident ISF proteoforms, we used to two methods to filter fragment proteoforms. First, fragment proteoforms were filtered based on their retention times because ISF proteoforms and the reference or alternative reference proteoform typically exhibit highly similar retention times. Second, fragment proteoforms were filtered based on the signal quality in MS1 spectra and the quality of the match between the proteoform and its MS/MS spectrum. Specifically, in an MS data file, a proteoform was selected as an ISF proteoform if (1) its apex retention time was within 6 seconds from the reference elution time, (2) its ECScore, a confidence score reported by TopFD [37], was at least 0.5, and (3) the proteoform was matched to an MS/MS spectrum with an E-value ≤ 0.01, which was reported by TopPIC [38].

For each MS data file, we calculated the relative intensity of each proteoform within the ISF proteoform group as the ratio between the signal intensity of the proteoform and the total signal intensity of all proteoforms in the ISF proteoform group. The relative intensity is referred to as the *ISF relative intensity (ISF-RI)* of the proteoform in the data file. To calculate the average ISF-RI across technical triplicates, only replicates in which the proteoform was detected were included. The *apex ISF-RI* of a proteoform was defined as the highest ISF-RI value, averaged across technical triplicates, observed among the 11 ISF voltage settings.

### 2.4 Middle-down MS

Myoglobin and CA2 were analyzed using middle-down MS. A total of 50 μg of protein was dissolved in 100 μL of 50 mM ABC with 8 M urea (pH 8.0). The protein solution was then reduced with 1 μL of 1 M DTT at 37 °C for 30 minutes and alkylated with 2.5 μL of 1 M IAA at 23 °C for 30 minutes. The resulting protein was digested separately with five enzymes: AspN, LysC, GluC, chymotrypsin, and trypsin, at 37 °C for 3 min using an enzyme-to-protein ratio of 1:50 (w/w). After digestion, the solution was acidified with 100 μL of TFA to a final concentration of 0.5% (v/v) to terminate the reaction. The resulting peptides/proteoforms were desalted using a C18 cartridge column (Thermo Scientific, Marietta, OH), followed by lyophilization in a vacuum concentrator (Thermo Fisher Scientific, Marietta, OH). The dried samples were resuspended in 50 μL of water containing 0.1% FA, quantified using Pierce Protein Assay (Thermo Fisher Scientific, Marietta, OH), and stored at −20 °C until use.

A total of 100 ng of peptides/proteoforms were separated by RPLC using a C2 column (300 Å, 3 μm, 100 μm i.d., 60 cm length, CoAnn, Richland, WA). In the RPLC system, the mobile phases were the same as those used in the top-down MS experiments. A 45-minute gradient for phase B was applied as follows: 0-2 min, 5% to 35%; 2-5 min, 35% to 50%; and 5-45 min, 50% to 80%. For each digested sample, triplicate MS runs were performed using CID and HCD fragmentation (3 runs for CID and 3 runs for HCD). MS1 and MS/MS spectra were collected using the same Orbitrap Fusion Lumos mass spectrometer and the same settings as the top-down MS analysis except for the settings of the NCE, which was set to 40% for CID runs and set to 35% for HCD runs.

### 2.5 Middle-down MS data analysis

In middle-down MS data preprocessing, msconvert [36] was used for converting raw data into centroided mzML files. Only spectra with retention times between 0 - 75 minutes were kept because 75 minutes was the total time for the programmed gradient for peptide/proteoform separation and an additional dwell time caused by the delay in the LC system. TopFD (version 1.7.9 and parameter settings in Supplemental **Tables S1**) was used for spectral deconvolution and peptide/proteoform feature detection. Then the deconvoluted spectra were searched against a database containing only the target protein sequence for peptide/proteoform identification using TopPIC (version 1.7.9 and parameter settings in Supplemental **Tables S2**). Peptide/proteoform identifications reported by TopPIC were further filtered using the confidence score of its peptide/proteoform feature: A peptide/proteoform identification was removed if the ECScore of its feature is less than 0.5.

Given an MS data file and a group of peptide/proteoform identifications, the relative intensity (RI) of a peptide/proteoform with respect to the group was calculated as the ratio of its intensity to the total intensity of all peptides/proteoforms in the group. To compare the abundances of peptides/proteoforms with various lengths, we also normalized peptide/proteoform intensities by their lengths. The normalized intensity of a peptide/proteoform with *L* amino acids and a feature intensity *I* is defined as *I* × *L*. The normalized relative intensity (NRI) of the peptide/proteoform is the ratio of its normalized relative intensity to the total normalized relative intensity of all peptide/proteoform identifications in the group.

The middle-down MS raw files were also searched against a database containing only the target protein sequence using the closed search mode of MSFragger [39] (version 23.1 and parameter settings in Supplemental **Table S3**). The mass tolerances for both precursor and fragment masses were set to 10 ppm. Acetylation at the protein N-terminus was specified as a variable PTM, and carbamidomethylation on cysteine was set as the fixed PTM. For each enzyme, we determined the largest number of missed cleavage sites in fragment proteoforms/peptides reported by TopPIC from the MS files and set this value as the maximum number of missed cleavage sites in MSFragger. Peptide-spectrum-match (PSM) identifications were filtered using an E-value cutoff of 0.01.

## 3 Results

### 3.1 In-source proteoform fragmentation

We evaluated in-source proteoform fragmentation using three proteins: ubiquitin (8,559.62 Da), myoglobin (16,940.97 Da), and CA2 (29,006.68 Da) by conducted 11 top-down LC-MS runs for each of the three proteins varying the ISF energy from 0 V to 100 V in a 10 V increment (see **Methods**).

#### 3.1.1 Reference, alternative reference, and ISF proteoforms

The most abundant proteoform in the LC-MS run with an ISF energy of 0V was selected as the reference proteoform. Proteoforms identified from the TD-MS files were divided into two groups: full-length proteoforms and fragment proteoforms. A proteoform containing all amino acids or all amino acids except for the N-terminal methionine of a protein is a full-length proteoform. Other proteoforms are fragment ones. The selected reference proteoform was a full-length proteoform with all amino acids for ubiquitin, a full-length proteoform with an N-terminal methionine excision (NME) for myoglobin, a full-length proteoform with an NME and an N-terminal acetylation for CA2 (Supplemental **Table S4**).

For myoglobin, the reference proteoform was not detected in the MS data with an ISF voltage of 100V. The most abundant proteoform identified in the MS file was a fragment proteoform corresponding to amino acid residues 101-154 with a mass of 5970.09 Da (Supplemental **Table S4**). Both the reference proteoform and the fragment proteoform were detected in the MS data with an ISF voltage of 80V. Because their extracted-ion chromatograms (XICs) were highly similar (Supplemental **Fig. S1(a)**), the fragment proteoform was selected as the alternative reference proteoform for myoglobin for the MS data with an ISF voltage of 100V.

For CA2, the reference proteoform was not detected in the MS data with ISF voltages of 70V and above. The most abundant proteoform in the MS data with an ISF voltage of 70V was a fragment proteoform of amino acid residues 200-260 with a mass of 7040.79 Da (Supplemental **Table S4**), which was also observed in the MS data files with an ISF voltage of 80V, 90V, and 100V. Because their XICs were highly similar (Supplemental **Fig. S1(b)**) in the MS data file with an ISF voltage of 50V, the fragment proteoform was selected as an alternative reference for MS data files of CA2 with an ISF voltage of 70V and above.

In addition to the reference proteoforms, we also identified other full-length proteoforms for the three proteins in the LC-MS run with an ISF energy of 0V. Specifically, four additional full-length proteoforms were detected for ubiquitin (with a single oxidation, two oxidations, water loss, or N-terminal acetylation), five for myoglobin (with a single oxidation, two oxidations, water loss, N-terminal methionine retention, or N-terminal acetylation), and three for CA2 (with a single oxidation, two oxidations, or water loss) (Supplemental **Table S5**). As the protein sample contained several full-length proteoforms, the ISF proteoforms observed in an MS file were a mixture of the products of several proteoforms.

We investigated if different full-length proteoforms can be separated by their retention times. We compared the apex retention times (ARTs) of the reference proteoform and other full-length proteoforms of the three proteins. Compared with the reference proteoform, the ARTs of the oxidized proteoforms were 7.7 - 14.9 seconds shorter for myoglobin, 13.9 - 21.4 seconds shorter for ubiquitin, and 10.4 - 59.9 seconds shorter for CA2. The ARTs of the proteoforms with N-terminal acetylation was 10.1 – 26.5 seconds longer than the reference proteoform for ubiquitin and myoglobin. The ARTs of the proteoforms with N-terminal methionine retention were even longer compared with the reference proteoform of myoglobin, with ART differences of 90.7 - 140.4 seconds. Because proteoforms with water loss are frequently generated during ionization, their ARTs typically cannot be distinguished from those of the corresponding proteoforms without water loss. The reference proteoform and its ISF proteoforms generally exhibit similar ARTs; however, the comparison results showed that the reference proteoform and other full-length proteoforms may also share similar ARTs. Therefore, using solely ART differences (e.g., a cutoff of 6 seconds) can only partially distinguish the ISF proteoforms of the reference proteoform from those of other full-length or intact proteoforms.

The reference proteoform and all selected ISF proteoforms across the 11 ISF voltages were combined to form the ISF proteoform group for each protein. The sizes of the ISF proteoform groups were 26 for ubiquitin, 25 for myoglobin, and 28 for CA2 (Supplemental **Tables S6–S8**). These ISF proteoform groups contained 8, 3, and 10 proteoforms covering the N-terminus of ubiquitin, myoglobin, and CA2, respectively. All N-terminal proteoforms shared the same N-terminal form as their corresponding reference proteoform, suggesting that ART-based filtering may have removed most ISF proteoforms originating from other intact proteoforms.

#### 3.1.2 Abundances of reference and ISF proteoforms

We assessed the relationship between ISF proteoforms and collision energy settings. A consistent trend was observed across all three proteins: the majority of ISF proteoforms, especially shorter ones, were observed at ISF energies above 50 V (Supplemental **Fig. S2**). We then examined the abundances of reference proteoforms and their ISF proteoforms across 11 ISF energy settings. For each protein, we selected three representative ISF proteoforms (Supplemental **Table S9**) and compared the ISF-RIs (see **Methods**) of the reference proteoform with those of the three representative ISF proteoforms (**Fig. 1**). The ISF-RIs of the reference proteoforms of all three proteins decreased as the ISF voltage increased. At lower ISF energy settings, the ISF-RIs of the reference proteoforms remained high. For ubiquitin, the ISF-RI of the reference proteoform decreased from over 95% to less than 15% as the ISF energy increased from 0 V to 100 V. A similar trend was observed for myoglobin and CA2. The ISF-RI of the reference proteoform of CA2 declined more rapidly than those of ubiquitin and myoglobin, becoming undetectable after 70 V. A possible explanation is that larger proteoforms tend to be fragmented more easily by ISF than smaller ones.

**Fig. 1:**
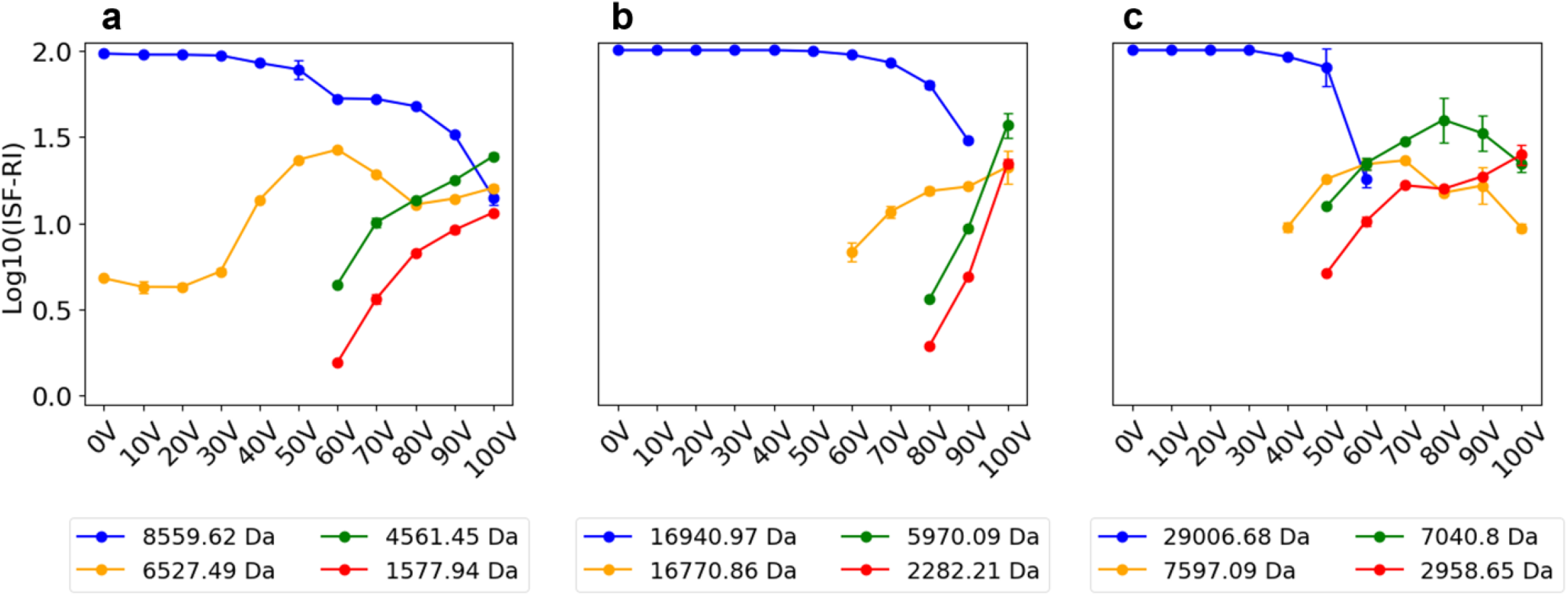
ISF-RIs of the reference proteoform and three representative ISF proteoforms across various ISF energy settings in top-down MS. The average ISF-RIs from triplicate MS runs for the four proteoforms of (a) ubiquitin, (b) myoglobin, and (c) CA2.

For ubiquitin, one representative proteoform with a mass of 6527.49 Da and a length of 58 (amino acids from 19 to 76) amino acids was observed in all runs with various ISF settings. Its ISF-RI increased as the ISF energy rose from 0V to 60V and decreased as the ISF energy continued to increase. Some shorter proteoforms, such as those with masses of 4561.45 Da (amino acids from 37 to 76) and 1577.94 Da (amino acids from 63 to 76), were observed only at high ISF energies, and both peaking at 100V (**Fig. 1a**). These results suggest that the abundances of ISF proteoforms are determined not only by the ISF energy but also by their amino acid sequences. Similar ISF-RI patterns of representative fragment proteoforms were observed for myoglobin and CA2 (**Fig. 1b, 1c**). At high ISF energies, the reference proteoforms became undetectable. In myoglobin, the reference proteoform was absent from the MS data at 100 V, where the most abundant proteoform was a fragment of 5970.09 Da corresponding to amino acids 101–154. This fragment proteoform first appeared at 80 V, and its ISF-RI increased with higher ISF energy, reaching a maximum at 100 V.

For CA2, the reference proteoform was not detected in the MS data at ISF energies of 70 V and above. The most abundant proteoform at 70 V was a fragment with a mass of 7040.09 Da, corresponding to amino acids 200–260. The fragment proteoform was first observed at 40 V, reached its apex ISF-RI at 70 V, and was also present at 80 V, 90 V, and 100 V. These observations suggest that as ISF energy increases, large proteoforms tend to be fragmented into smaller species, and these small fragment proteoforms became dominant.

#### 3.1.3 Sequence coverage and ISF energy

We examined the sequence coverage of the three proteins with single ISF voltage settings. For a proteoform, a charge state, and an MS data file, the *representative proteoform-spectrum-match (PrSM)* of the proteoform in the file is the PrSM with a matched precursor charge state and the lowest E-value. For each MS file, representative PrSMs across all charge states were combined for each proteoform in the ISF proteoform group to enhance sequence coverage (Supplemental **Fig. S3**). The highest sequence coverage (92.4% for ubiquitin, 57.7% for myoglobin, and 39.2% for CA2) was achieved at ISF energies of 70V for ubiquitin, 80V for myoglobin, and 60V for CA2 (**Fig. 2a-c** and supplemental **Fig. S4**). The results showed that the single MS runs at the optimal ISF voltage yielded higher sequence coverage than the single MS runs at 0 V, although the improvement was modest.

**Fig. 2:**
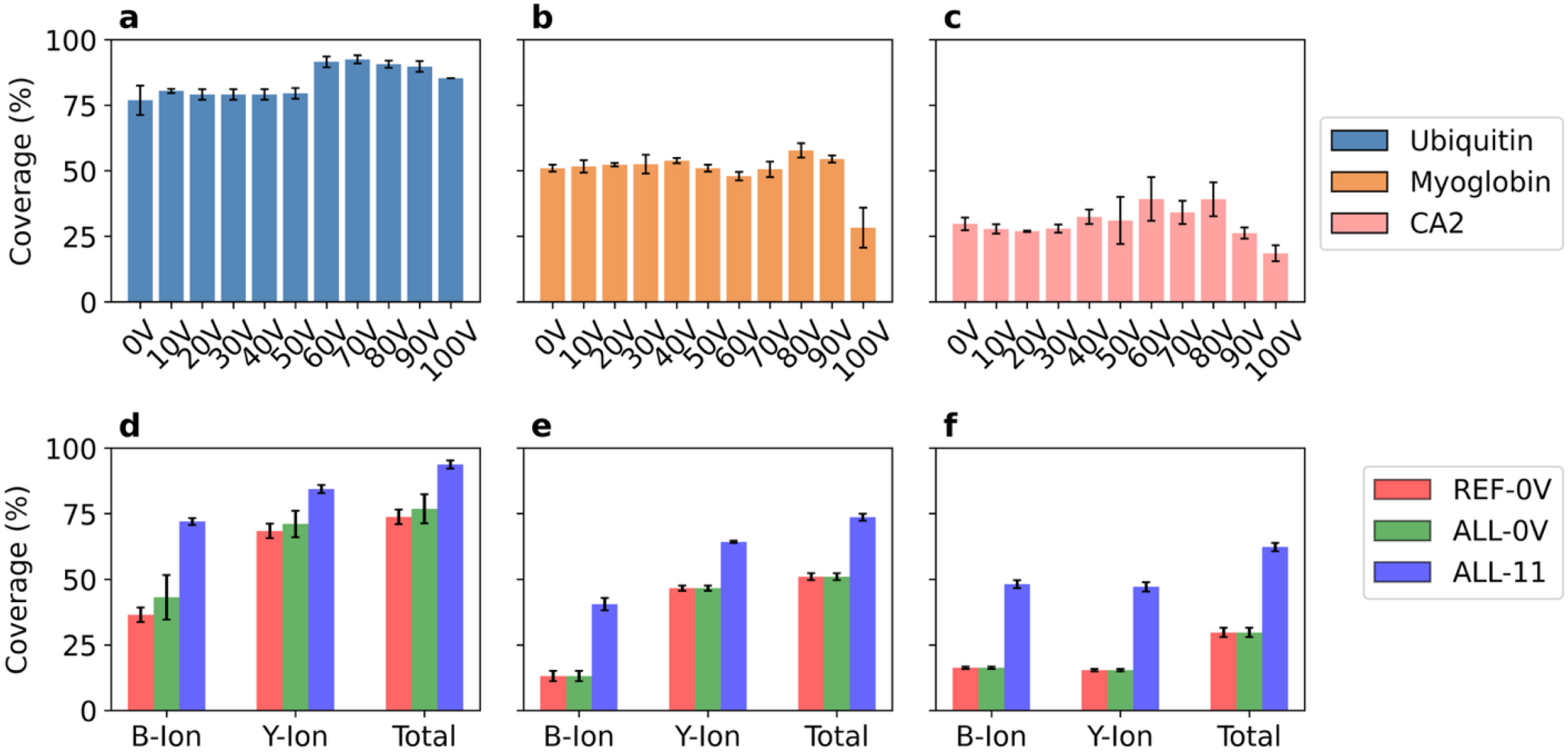
Sequence coverage obtained by ISF proteoforms across various ISF settings. The average sequence coverage obtained with various ISF settings in the triplicate MS runs for (a) ubiquitin, (b) myoglobin, and (c) CA2. The average protein sequence coverages obtained using the representative PrSMs of the reference proteoform at 0V (REF-0V), the representative PrSMs for each identified proteoform in the ISF group at 0V (ALL-0V), and the apex representative PrSMs of all ISF proteoforms in the ISF group in the 11 MS files (ALL-11) for (d) ubiquitin, (e) myoglobin, and (f) CA2. The error bars represent standard deviation.

We further evaluated whether combining top-down mass spectra obtained at various ISF energy settings could enhance fragment ion coverage of the protein sequences. Several short ISF proteoforms were observed only when the ISF energy exceeded 40 V for the three proteins, and the fragment ions from these proteoforms helped increase the fragment ion coverage. For each proteoform, we chose the MS file with highest sequence coverage as the apex MS file of the proteoform and selected the representative PrSMs of the proteoform in the apex MS file as apex representative PrSMs of the proteoform. Then the apex representative PrSMs of each proteoform in the ISF proteoform group were combined to compute the sequence coverage of a protein.

For each of the three proteins, we compared the fragment ion sequence coverage obtained using three approaches: (A) sequence coverage obtained by combining the representative PrSMs across all charge states of the reference proteoform at the ISF energy of 0V, (B) sequence coverage obtained by combining the representative PrSMs across all charge states of all proteoforms in the ISF proteoform group identified at the ISF energy of 0V, and (C) coverage obtained by combining the apex representative PrSMs across all charge states of all ISF proteoforms. The proteoform sequence coverage of ubiquitin increased from 73.8% with method A to 76.9% with method B, and further to 93.8% with method C **(Fig. 2d)**. Method C also achieved sequence coverages of 73.6% for myoglobin and 62.3% for CA2, representing substantial improvements compared with the other two methods (**Fig. 2e-f**).

### 3.2 Middle-down proteomics

#### 3.2.1 Digestion efficiency of five enzymes in middle-down MS

We evaluated the digestion efficiency of five enzymes (AspN, chymotrypsin, GluC, LysC, and trypsin) in middle-down MS using a 3-minute digestion of myoglobin and CA2, which left a portion of intact proteoforms undigested. We observed six full-length proteoforms of myoglobin and four full-length proteoforms of CA2 (Supplemental **Table S10**), so the digested proteoforms/peptides were generated from a mixture of intact proteoforms. For an MS run, the digestion ratio (DR) was calculated as the total RI of all fragment proteoforms/peptides. Because a full-length proteoform may produce several digested proteoforms/peptides, the digestion ratio may overestimate the percentage of proteoforms that are digested. To address this problem, we also calculated the normalized digestion ratio (NDR) of an MS run as the total NRI of all fragment proteoforms/peptides (**Methods**). Because some digested proteoforms/peptides are unidentified, the NDR may be an underestimate of the percentage of proteoforms that are digested.

We compared the numbers of cleavage sites of the five enzymes in the two proteins (**Fig. 3a-b**). Chymotrypsin has the highest numbers of cleavage sites due to its broad substrate specificity, while AspN has the lowest numbers [36]. The number and distribution of the cleavage sites affect the length of digested proteoforms/peptides and digestion efficiency. We assessed the digestion efficiency of the 5 enzymes using DRs and NDRs in triplicate MS runs (**Fig. 3c-f**). GluC exhibited the highest digestion efficiency with an averaged DR of 49.6% and NDR of 34.0% for myoglobin, and both chymotrypsin and GluC achieved high digestion efficiency for CA2. LysC and trypsin showed intermediate efficiency, though consistently lower than those of chymotrypsin and GluC. Chymotrypsin contains the highest numbers of cleavage sites (30 for myoglobin and 55 for CA2) than other enzymes (**Fig. 3a-b**), which might contribute to its high digestion efficiency. While GluC, LysC, and trypsin have similar numbers of cleavage sites, a possible reason for GluC’s high digestion efficiency was that GluC had lower missed cleavage rate than the other enzymes. AspN consistently demonstrated the lowest digestion efficiency with a ∼1% DR for both proteins, which was the results of a small number of cleavage sites and a high missed cleavage rate. Notably, the DRs and NDRs of CA2 were consistently higher than myoglobin across all enzymes. The reason might be that the long sequence of CA2 provided more cleavage sites for digestion than myoglobin.

**Fig. 3:**
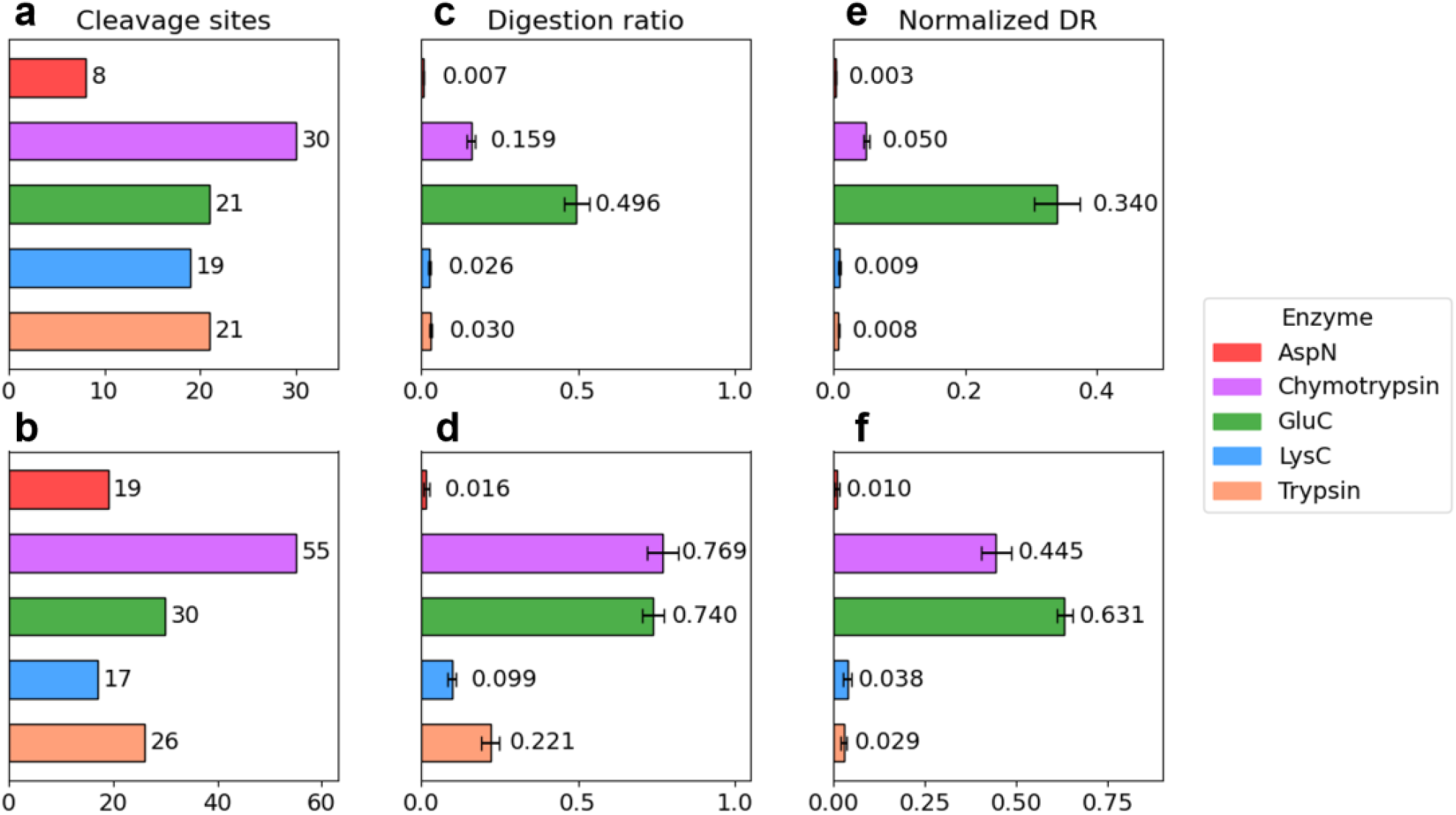
Digestion efficiency of five enzymes in 3-minute digestion on myoglobin and CA2. Numbers of potential cleavage sites in myoglobin (a) and CA2 (b). DRs of (c) myoglobin and (d) CA2 and NDRs of (e) myoglobin and (f) CA2 in the triplicate MS runs.

#### 3.2.2 Digested proteoforms and peptides

For myoglobin, chymotrypsin, GluC, and trypsin each generated approximately 50 peptides/proteoforms, while AspN and LysC produced about 35. For CA2, chymotrypsin and trypsin yielded substantially more peptides/proteoforms than the other enzymes (**Fig. 4a-b**)

**Fig. 4:**
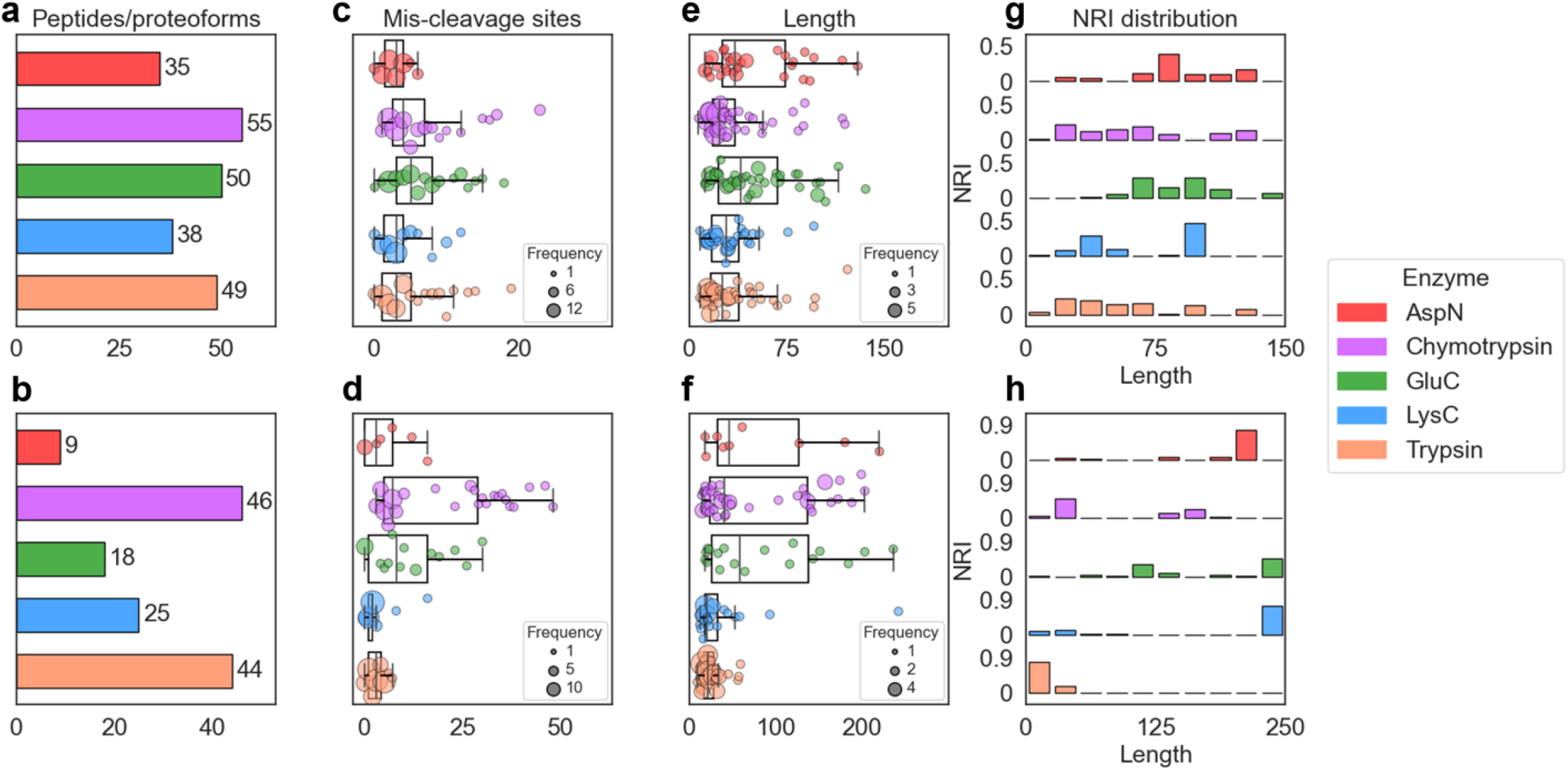
Comparison of digested proteoform/peptides of myoglobin and CA2 in MD-MS using five enzymes. The proteoforms/peptides generated by the five enzymes in the first replicate of the MD-MS runs were studied. Numbers of digested proteoform/peptides for myoglobin (a) and CA2 (b); distributions of MCSs of digested proteoforms/peptides for myoglobin (c) and CA2 (d); distributions of proteoform/peptide lengths for myoglobin (e) and CA2 (f); and histograms of normalized relative intensities versus proteoform/peptide lengths for myoglobin (g) and CA2 (h).

We examined the numbers of missed cleavage sites (MCSs) in the digested proteoforms/peptides produced by each enzyme **(Fig. 4c-d** and supplemental **Fig. S5, S6)**. Chymotrypsin and GluC had the largest numbers of cleavage sites in the two proteins (**Fig. 3a-b**), and their digested peptides/proteoforms contained more MCSs than the other enzymes. Some digested proteoforms/peptides of chymotrypsin and GluC had 20-40 MCSs, showing their high missed cleavage rates in the 3-min digestion. While the average numbers of MCSs of GluC were slightly higher than chymotrypsin, the variances of MCSs of chymotrypsin were slightly higher than GluC. The digested proteoforms/peptides of the other three enzymes contained fewer than 5 MCSs in myoglobin and fewer than 8 MCSs in CA2 on average, which was much higher than the common 0 or 1 MCS observed in BUP, showing that the 3-min digestion significantly increased the missed cleavage rate for these enzymes compared with long digestion time in BUP.

We also studied the length of the digested proteoforms/peptides for each enzyme **(Fig. 4e-f** and supplemental **Fig. S5, S6)**. Both GluC and AspN generated many long digested proteoforms/peptides, but the reasons were different: the main reason for GluC was its high missed cleavage rate and that for AspN was its few cleavage sites. While chymotrypsin, LysC, and trypsin produced mainly short peptides with less than 50 amino acids, different patterns were observed between myoglobin and CA2. The average lengths of digested proteoforms/peptides of myoglobin were similar for the three enzymes, but LysC and trypsin produced shorter peptides than chymotrypsin for CA2, suggesting the missed cleavage rate was affected the amino acid sequence of the protein. Among the five enzymes, AspN yielded the fewest digested proteoforms/peptides due to its limited number of cleavage sites.

We also studied the abundances of the digested proteoforms/peptides produced by the five enzymes (**Fig. 4g-h**). For myoglobin, high abundance proteoforms/peptides were mainly long ones for AspN and GluC (>75 amino acids) and mainly short ones for LysC and trypsin except that LysC had a high abundance proteoform with a length of 96. The abundance distribution for chymotrypsin was balanced between long and short ones. For CA2, the highest abundance proteoforms were long ones for AspN, GluC, and LysC. Specifically, the dominate proteoforms had a length of 220 amino acids, 236 amino acids, 242 amino acids for AspN, GluC and LysC, respectively. The proteoforms for trypsin were dominated by short ones.

#### 3.2.3 Sequence coverage in MD-MS

While the TopFD [37] is suited for deconvoluted large fragment masses in MD MS/MS spectra, it sometimes misses small fragment masses in spectral deconvolution. Database search software tools for BUP, such as MSFragger [39], match peptides to MS/MS spectra without spectral deconvolution, making them efficient to match small fragment masses to peptides. Because of this, we combined PrSMs reported by TopPIC and peptide-spectrum-matches (PSMs) reported by MSFragger to increase sequence coverage.

We examined the sequence coverage of the two proteins obtained by MD-MS with the five enzymes. For myoglobin, GluC and chymotrypsin achieved the highest sequence coverage (**Fig. 5a-b**), likely due to their high DRs (**Fig. 3c**) and their ability to produce many long digested proteoforms (**Fig. 4e**). Although AspN also produced long digested proteoforms (**Fig. 4e**), it generated only on average 34.7 digested proteoforms/peptides (Supplemental **Fig. S5**), limiting its sequence coverage. Chymotrypsin, LysC, and trypsin provided intermediate coverages for myoglobin. Combining proteoforms/peptides generated by the five enzymes resulted in near complete sequence coverage for myoglobin: 99.3% with CID and 98.7% with HCD.

**Fig. 5:**
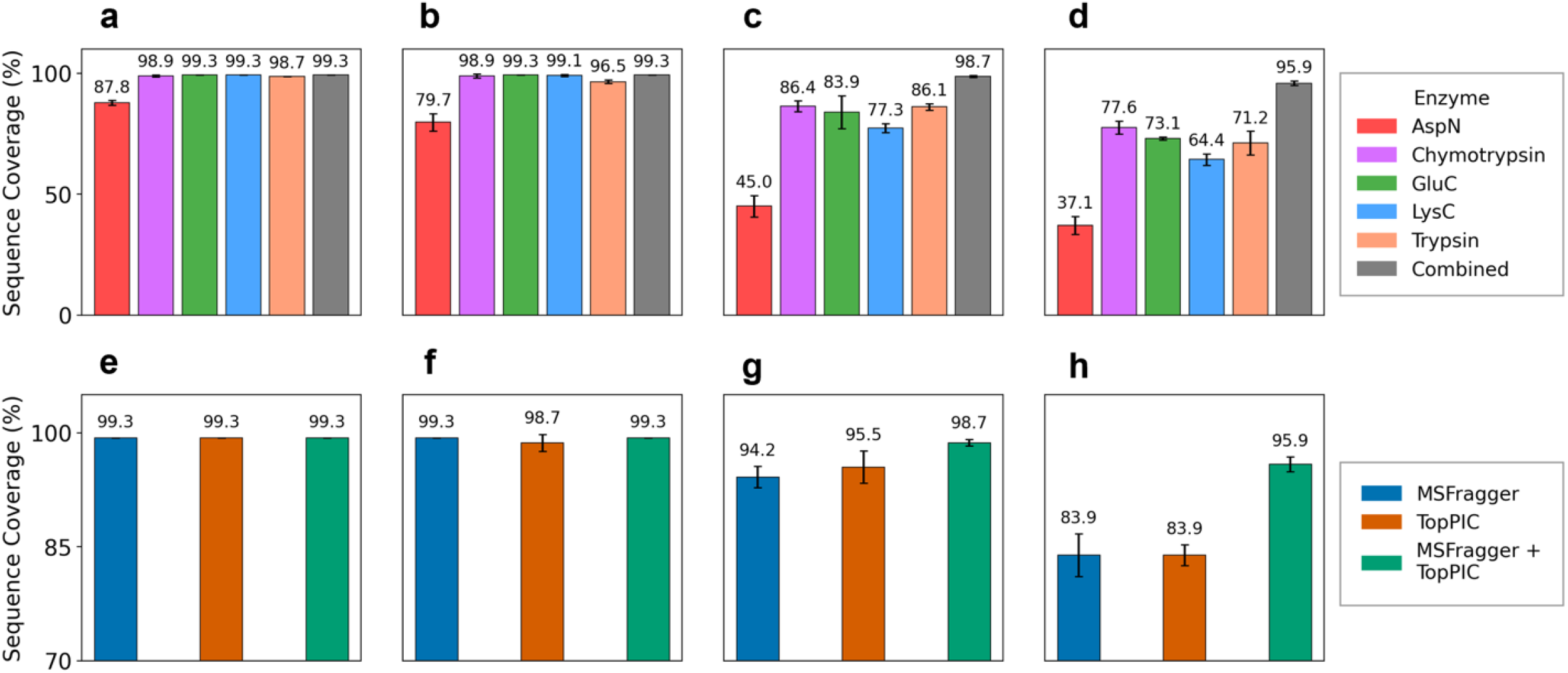
Sequence coverage of myoglobin and CA2 in MD-MS using five enzymes. Sequence coverage comparison across five enzymes: (a) myoglobin with CID, (b) myoglobin with HCD, (c) CA2 with CID, and (d) CA2 with HCD. Sequence coverage comparison among the MSFragger-only method, the TopPIC-only method, and the combined method using data from all five enzymes: (e) myoglobin with CID, (f) myoglobin with HCD, (g) CA2 with CID, and (h) CA2 with HCD.

Similarly, chymotrypsin and GluC yielded the highest sequence coverage and AspN reported the lowest sequence coverage among the five enzymes for CA2 (**Fig. 5c-d**). And combining proteoforms/peptides generated by the five enzymes resulted in a high sequence coverage of 95.5% with CID and 83.9% with HCD. For single enzymes, the sequence coverages of CA2 were lower than myoglobin. A possible reason is that CA2 is longer than myoglobin, making it more challenging to obtain high sequence coverage. In addition, CID consistently provided higher sequence coverage than HCD across all enzymes (**Fig. 5a-d**).

We also compared the sequence coverages obtained using three types of data: (A) PrSMs reported by TopPIC, (B) PSMs reported by MSFragger, and (C) PrSMs reported by TopPIC and PSMs reported by MSFragger. Compared with the TopPIC only method, the combined method increased the sequence coverage slightly for myoglobin and significantly for CA2 (**Fig. 5e-h**). Further examination of the matched fragment masses identified by TopPIC and MSFragger revealed that PSMs reported by MSFragger provided many matched small fragment masses that were missed by spectral deconvolution in the TopFD–TopPIC analysis pipeline (Supplemental **Fig. S7-S11**).

## 4 Discussion and Conclusions

In this paper, we evaluated ISF in TD-MS and short-time enzymatic digestion in MD-MS to enhance proteoform sequence coverage using three proteins: ubiquitin, myoglobin, and CA2. Pseudo-MS^3^ spectra generated by TD-MS with ISF improved fragment ion coverage for all three proteins compared with TD-MS without ISF, most notably for ubiquitin, which achieved sequence coverage exceeding 90% **(Fig. 2a and 2d)**. However, using TD-MS with ISF failed to achieve almost complete sequence coverage for myoglobin and CA2, possibly due to the limited numbers of fragment proteoforms generated by ISF. For myoglobin and CA2, the reference proteoforms became undetectable at high ISF energies **(Fig. 1b and 1c)**, suggesting that longer proteoforms may be more easily fragmented by ISF than shorter ones.

MD-MS experiments with myoglobin and CA2 showed that a short enzymatic digestion time (3 minutes) left some intact proteoforms undigested and produced a mixture of digested long proteoforms and short peptides. Combining the peptides and proteoforms generated by the five enzymes achieved more than 90% sequence coverage for both proteins, demonstrating that this approach provides rich fragment information for PTM characterization and protein *de novo* sequencing. While BUP with multiple enzyme digestions can also achieve high sequence coverage [40], the long proteoforms produced by MD-MS provide additional information on PTM combinations. Because TDP, MDP, and BUP have complementary strengths in proteoform characterization, integrating the three strategies can yield better sequence coverage than any single approach.

The results from MD-MS also highlight enzyme-specific differences in digestion efficiency and digested peptides/proteoforms. Among the enzymes tested, GluC and chymotrypsin achieved the highest protein sequence coverage. These findings suggest that enzyme selection is critical for optimizing MD-MS workflows for achieving high sequence coverage.

This study has several limitations. First, proteoforms are often coeluted in TD-MS analysis of complex samples, making it challenging to assign ISF proteoforms generated from coeluted species to their corresponding intact proteoforms. Improved separation methods are needed to address this issue. Second, the analysis focused only on proteoforms smaller than 30 kDa. In the future, we will use these methods to study larger and more complex proteoforms.

## Data availability

The MS raw data can be downloaded from the PRIDE repository with the data set identifier

**PXD068831**.

## Acknowledgements

This research was funded by NSF through the grants 2307571, 2307572, and 2307573.

## Notes

The authors used ChatGPT to enhance the language and readability during the preparation of this paper. After utilizing ChatGPT, the authors reviewed and edited the content and take full responsibility for the final version of the paper.

