## Supplemental Table 1-3, Supplemental Figure 1-11 for "Enhancing proteoform sequence coverage using top-down mass spectrometry with in-source fragmentation and middle-down mass spectrometry"

**Table S1:** Parameters settings for TopFD (version 1.7.9)

| **Input Parameter** | **Value** |
|---|---|
| Maximum charge | 30 for ubiquitin and myoglobin, 40 for CA2 |
| Maximum mass | 40,000 Da |
| MS1 signal noise ratio | 3.0 |
| MS/MS signal noise ratio | 1.0 |
| *M/z* error tolerance | 0.02 |
| Min scan number in features | 3 |
| Use single scan noise level during feature extraction | True |
| ECScore cutoff | 0.5 |
| Fragmentation method | File |
| Disable additional feature search | True |

**Table S2:** Parameters settings for TopPIC (version 1.7.9)

| TopPIC Parameter | Value |
|---|---|
| **Proteome database** | Single protein sequence:<br>UniProt ID P0CH28 for ubiquitin<br>UniProt ID P68082 for myoglobin<br>UniProt ID P00921 for CA2 |
| **Search type** | Target |
| **N-terminal forms of proteins** | NONE, NME, M_ACETYLATION, NME_ACETYLATION |
| **Use TopFD features** | Yes |
| **Fixed modifications** | Top-down: None;<br>Middle-down: Carbamidomethylation on cysteine |
| **Variable PTM** | Top-down: Water loss, -18.01 Da, all the 20 amino acids, anywhere<br><br>Middle-down: None |
| **Maximum number of variable modifications** | 1 |
| **Spectrum level cutoff type for filtering PrSMs** | E-value |
| **The cutoff value for filtering PrSMs** | 0.01 |
| **Spectrum level cutoff type for filtering proteoforms** | E-value |
| **The cutoff value for filtering proteoforms** | 0.01 |
| **Error tolerance for precursor and fragment masses** | 10 ppm |
| **Error tolerance for identifying PrSM clusters** | 1.2 Da |
| **Maximum number of unexpected mass shifts** | 0 |
| **E-values computation** | Generating function |

**Table S3:** Parameters settings for MSFragger (version 23.1)

| **Input Parameter** | **Value** |
|---|---|
| Precursor mass tolerance (ppm) | 10 |
| Fragment mass tolerence (ppm) | 10 |
| Mass calibration | Enable |
| Isotope error | 0/1/2/3 |
| Cleavage | Selected based the enzyme used in the experiments: AspN, chymotrypsin, GluC, LysC, or trypsin |
| Missed cleavage number | Myoglobin AspN: 10 |
| | Myoglobin chymotrypsin: 30 |
| | Myoglobin GluC, LysC, trypsin: 20 |
| | CA2 AspN, LysC, trypsin: 20 |
| | CA2 chymotrypsin: 50 |
| | CA2 GluC: 30 |
| Peptide length (amino acids) | 6-100 |
| Peptide mass (Da) | 500-12000 |
| Variable PTMs | N-terminal acetylation |
| Fixed modification | Cysteine carbamidomethylation |
| Maximum variable modification on a peptide | 1 |
| Minimum peaks | 15 |
| Use top N peaks | 150 |
| Minimum ratio | 0.01 |
| Intensity transform | None |
| Remove precursor peak | Only peak with precursor charge |
| Report mass shift as a variable modification | No |
| Minimum matched fragments | 4 |
| Deisotope | Yes |
| Deneutralloss | Yes |

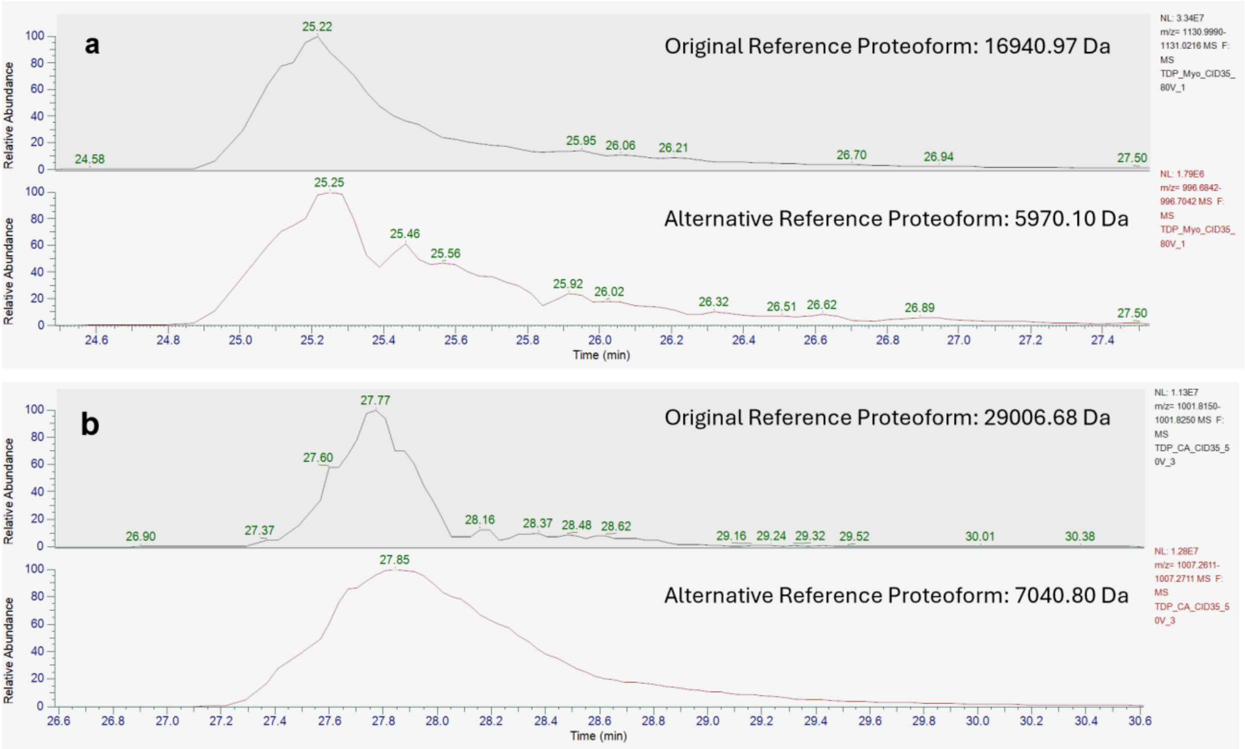


**Fig. S1: Comparison the XICs of the reference proteoform and alternative reference proteoform**. (a) An MS data file of myoglobin with an ISF voltage of 80 V and (b) an MS file of CA2 with an ISF voltage of 50 V.

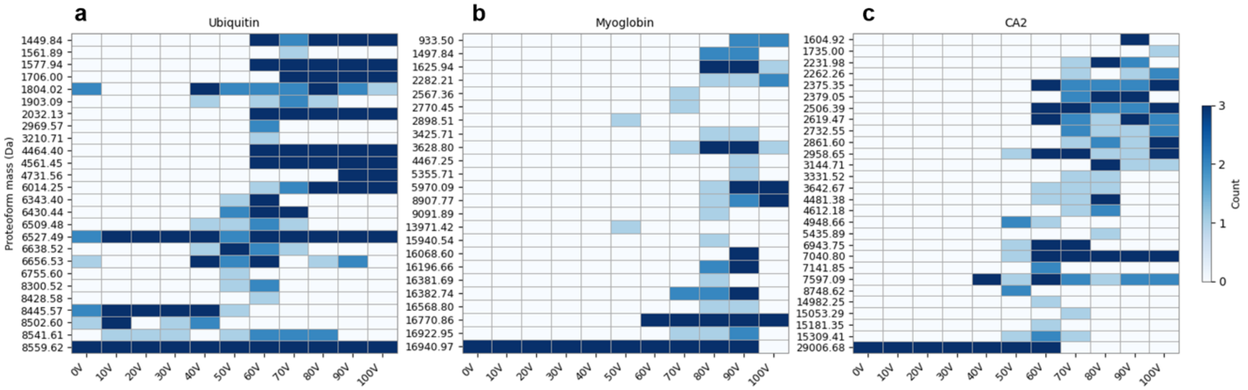


**Fig. S2: ISF proteoforms identified at different ISF voltage settings**. Proteoforms were ranked in the increasing order of their molecular masses. The color of each cell represents the number of technical MS replicates in which the proteoform was identified. (a) Ubiquitin, (b) myoglobin, and (c) CA2.

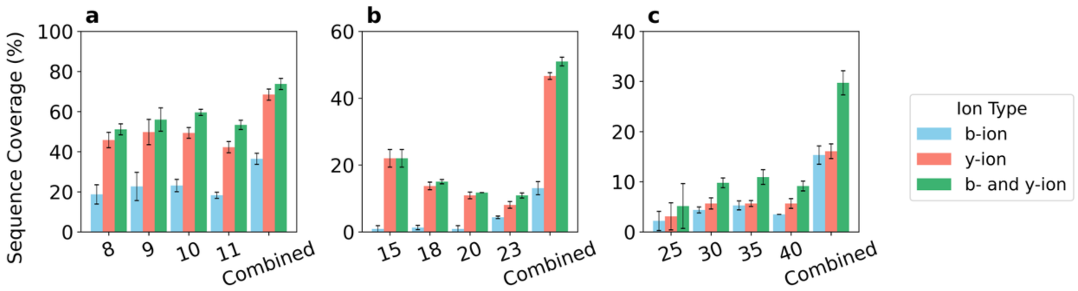


**Fig. S3: Comparison of proteoform sequence coverage using representative PrSMs of single charge states versus combining representative PrSMs of multiple charge states**. Sequence coverage obtained by representative PrSMs of the reference proteoform from triplicate MS runs at an ISF energy of 0 V are shown. (a) Single charge states 8, 9, 10, 11 are compared with combining multiple charge states 8 - 11 for ubiquitin. (b) Single charge states 15, 18, 20, 23 are compared with combining multiple charge states 15 - 23 for myoglobin. (c) Single charge states 26, 30, 35, 40 are compared with combining multiple charge states 25 - 40 for CA2.

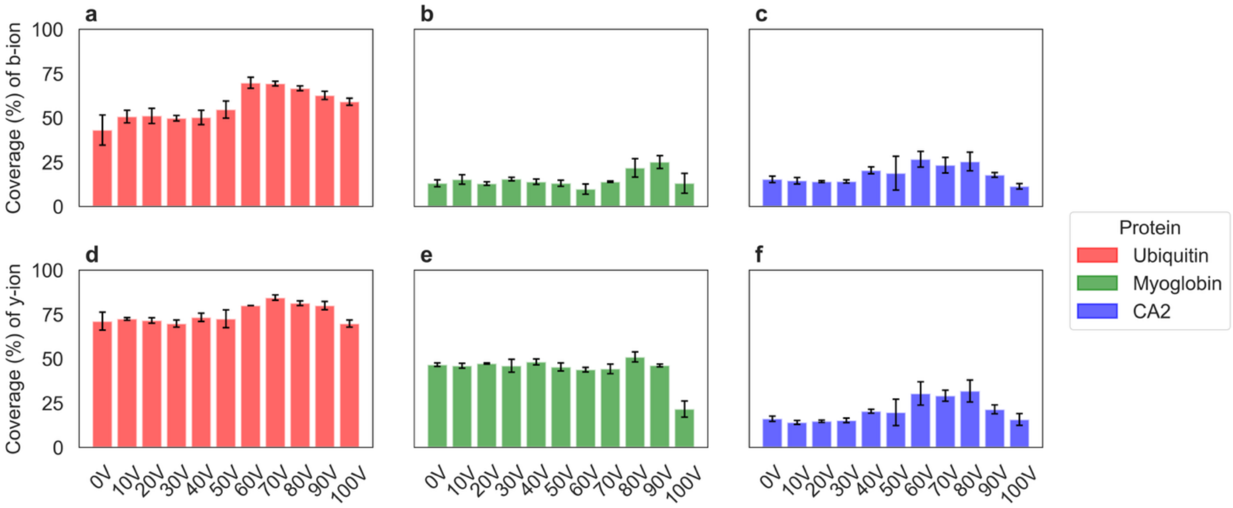


**Fig. S4: Comparison of proteoform sequence coverage from b-ions and y-ions across different ISF voltage settings.** B-ion coverage for (a) ubiquitin, (b) myoglobin, and (c) CA2, and y-ion coverage for (d) ubiquitin, (e) myoglobin, and (f) CA2. Error bars indicate standard deviations across triplicates.

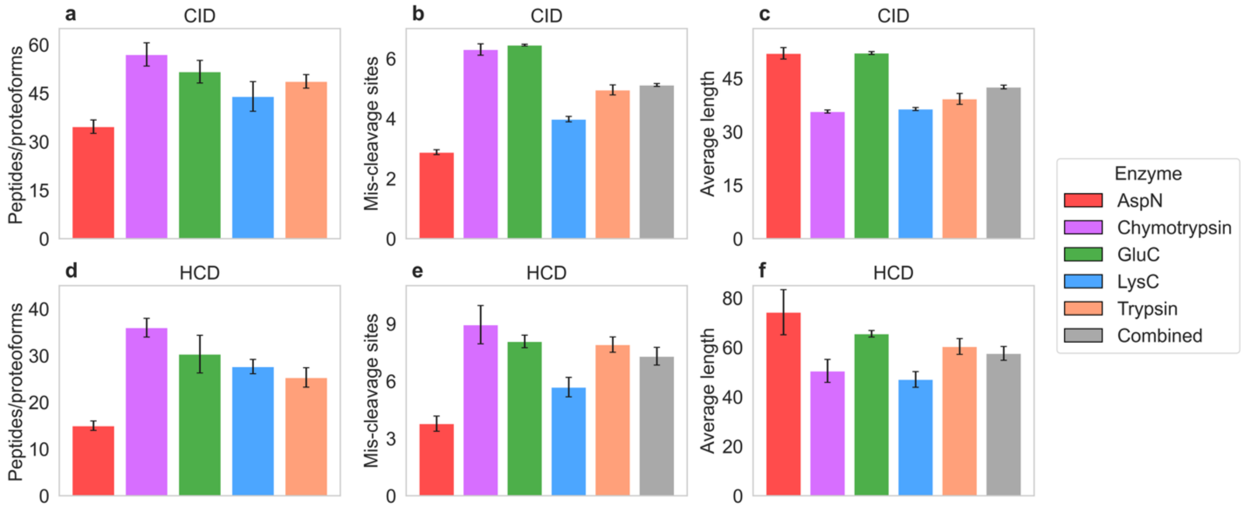


**Fig. S5: Comparison of digested peptides/proteoforms of myoglobin in MD-MS using five enzymes.** The numbers of digested peptides/proteoforms for CID (a) and HCD (d) runs; the average mis-cleavage sites in digested peptides/proteoforms for CID (b) and HCD (e) runs; and the average lengths of digested peptides/proteoforms for CID (c) and HCD (f) runs. Error bars represent standard deviations across triplicates.

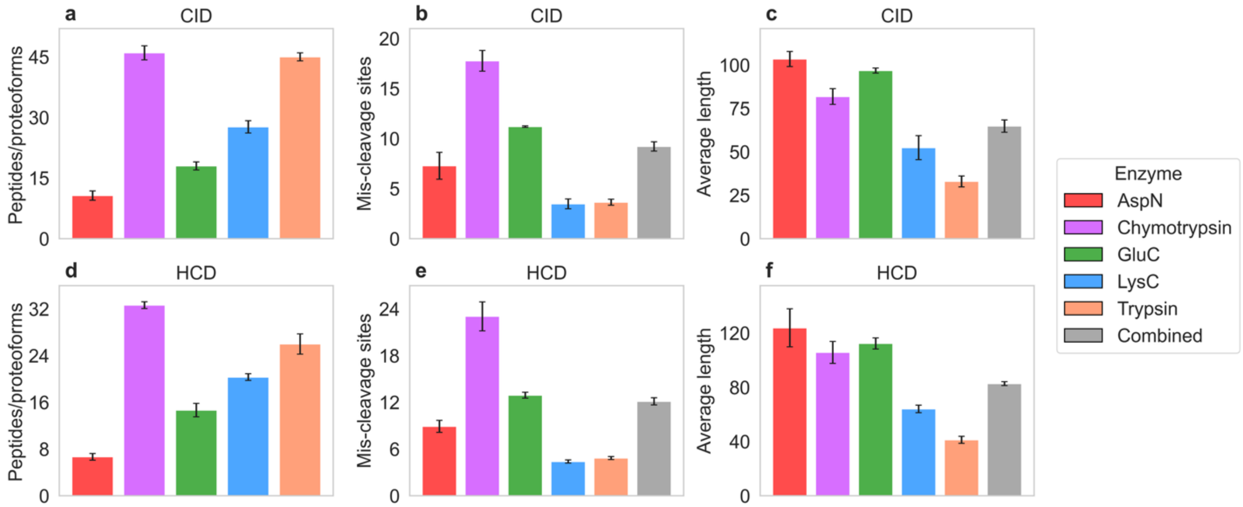


**Fig. S6: Comparison of digested peptides/proteoforms of CA2 in MD-MS using five enzymes.** The numbers of digested peptides/proteoforms for CID (a) and HCD (d) runs; the average mis-cleavage sites in digested peptides/proteoforms for CID (b) and HCD (e) runs; and the average lengths of digested peptides/proteoforms for CID (c) and HCD (f) runs. Error bars represent standard deviations across triplicates.

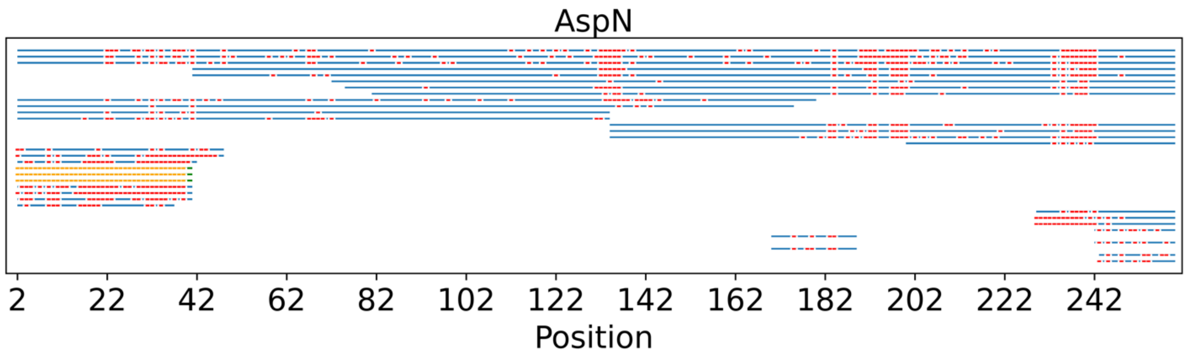


**Fig. S7: Peptide/proteoforms of CA2 identified by TopPIC and MSFragger in MD-MS using AspN digestion**. Identifications are from the first replicate of the CID runs. Each blue line with red ticks represents a peptide/proteoform identified by TopPIC, and each green line with orange ticks represents a peptide/proteoform identified by MSFragger. Red and orange ticks indicate matched fragment masses.

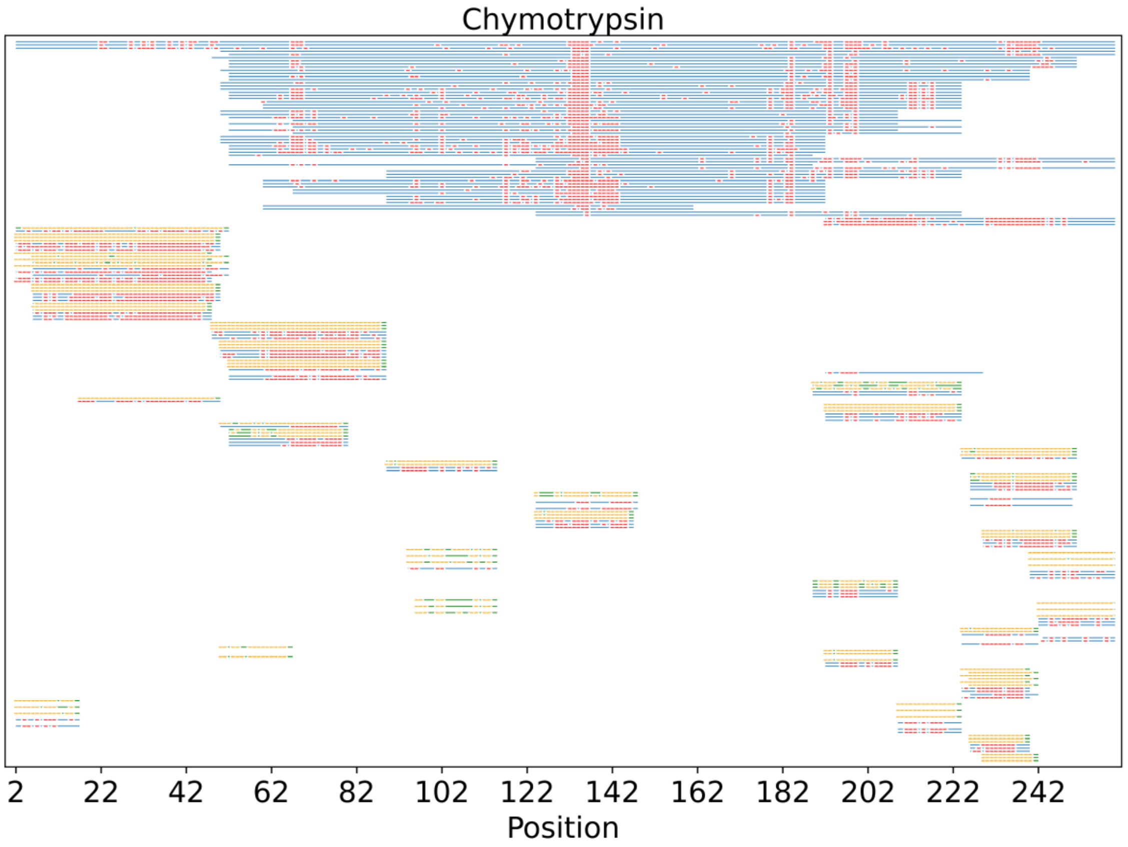


**Fig. S8: Peptide/proteoforms of CA2 identified by TopPIC and MSFragger in MD-MS using chymotrypsin digestion**. Identifications are from the first replicate of the CID runs. Each blue line with red ticks represents a peptide/proteoform identified by TopPIC, and each green line with orange ticks represents a peptide/proteoform identified by MSFragger. Red and orange ticks indicate matched fragment masses.

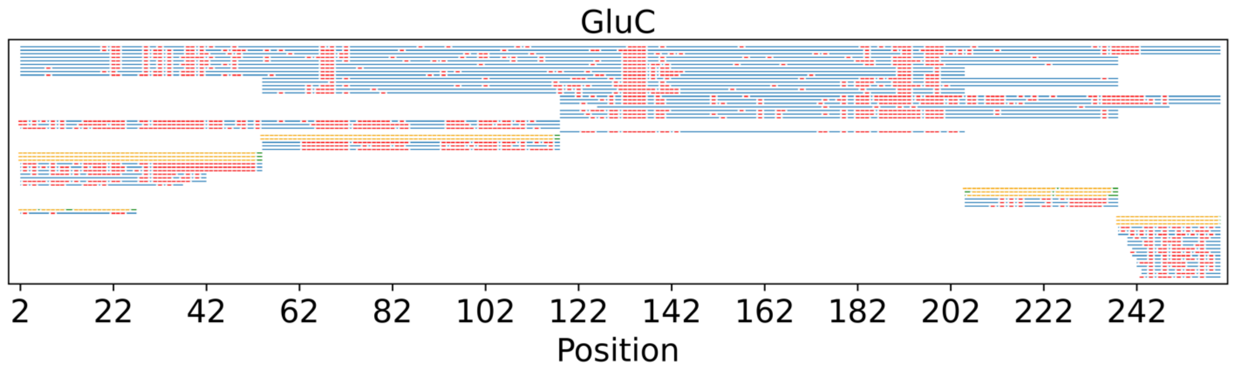


**Fig. S9: Peptide/proteoforms of CA2 identified by TopPIC and MSFragger in MD-MS using GluC digestion**. Identifications are from the first replicate of the CID runs. Each blue line with red ticks represents a peptide/proteoform identified by TopPIC, and each green line with orange ticks represents a peptide/proteoform identified by MSFragger. Red and orange ticks indicate matched fragment masses.

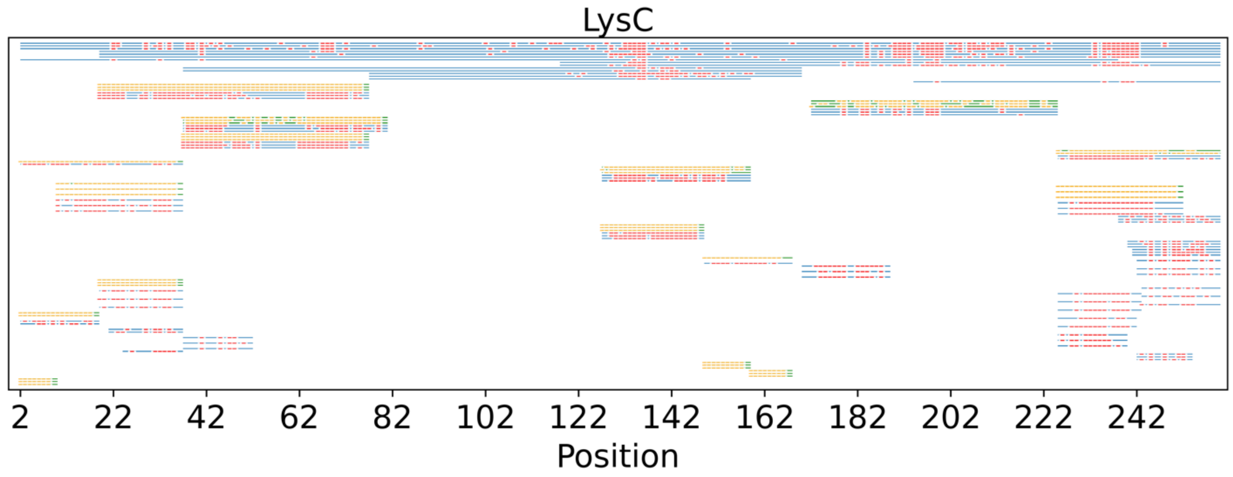


**Fig. S10: Peptide/proteoforms of CA2 identified by TopPIC and MSFragger in MD-MS using LysC digestion**. Identifications are from the first replicate of the CID runs. Each blue line with red ticks represents a peptide/proteoform identified by TopPIC, and each green line with orange ticks represents a peptide/proteoform identified by MSFragger. Red and orange ticks indicate matched fragment masses.

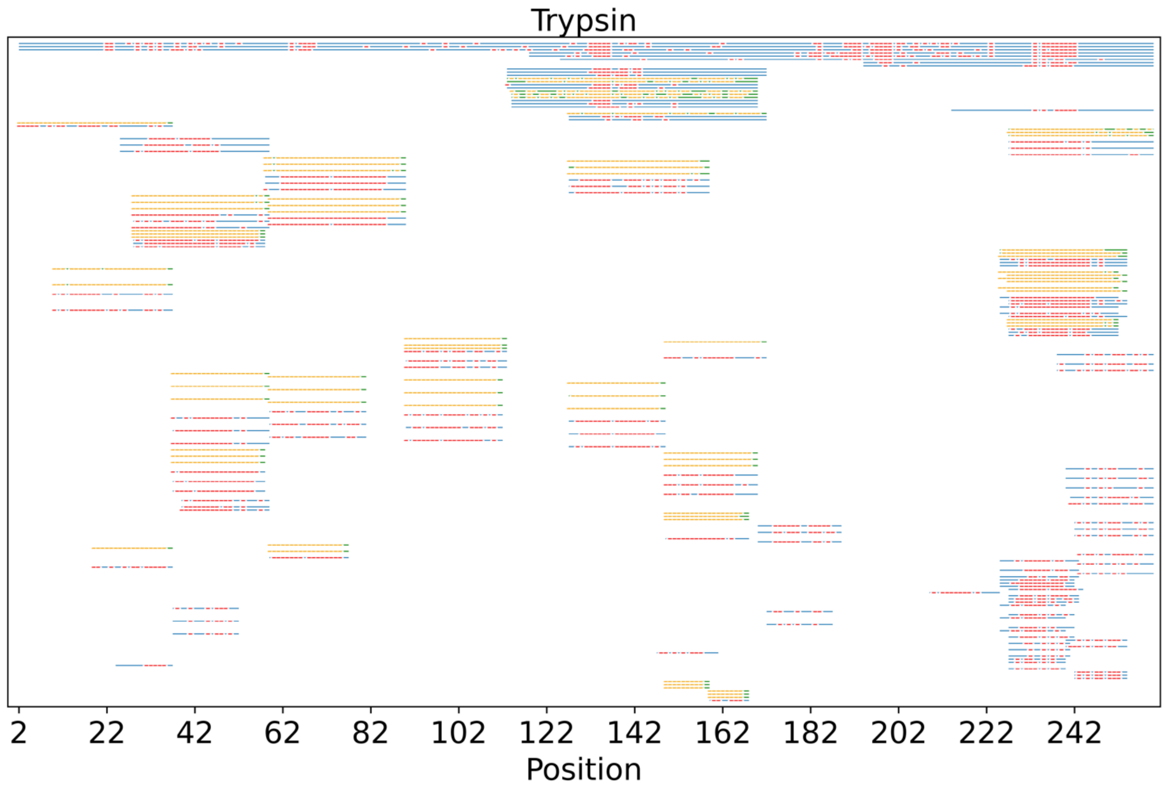


**Fig. S11: Peptide/proteoforms of CA2 identified by TopPIC and MSFragger in MD-MS using trypsin digestion**. Identifications are from the first replicate of the CID runs. Each blue line with red ticks represents a peptide/proteoform identified by TopPIC, and each green line with orange ticks represents a peptide/proteoform identified by MSFragger. Red and orange ticks indicate matched fragment masses.
